# Bayesian analysis of genetic association across tree-structured routine healthcare data in the UK Biobank

**DOI:** 10.1101/105122

**Authors:** Adrian Cortes, Calliope A. Dendrou, Allan Motyer, Luke Jostins, Damjan Vukcevic, Alexander Dilthey, Peter Donnelly, Stephen Leslie, Lars Fugger, Gil McVean

## Abstract

Genetic discovery from the multitude of phenotypes extractable from routine healthcare data has the ability to radically transform our understanding of the human phenome, thereby accelerating progress towards precision medicine. However, a critical question when analysing high-dimensional and heterogeneous data is how to interrogate increasingly specific subphenotypes whilst retaining statistical power to detect genetic associations. Here we develop and employ a novel Bayesian analysis framework that exploits the hierarchical structure of diagnosis classifications to jointly analyse genetic variants against UK Biobank healthcare phenotypes. Our method displays a more than 20% increase in power to detect genetic effects over other approaches, such that we uncover the broader burden of genetic variation: we identify associations with over 2,000 diagnostic terms. We find novel associations with common immune-mediated diseases (IMD), we reveal the extent of genetic sharing between specific IMDs, and we expose differences in disease perception or diagnosis with potential clinical implications.

---

Large-scale, hypothesis-free approaches for identifying genetic risk variants, including genome-wide association studies (GWAS) and next generation sequencing analyses, have greatly advanced our understanding of complex traits, with implications for drug development and clinical practice^1–5^. These approaches have typically involved genetic discovery from case-control cohorts where single clinically derived phenotypes are considered one at a time. However, resources such as the UK Biobank^6,7^, which has prospectively collected extensive health-relevant phenotypic and genotypic information from over 500,000 participants, allow for the simultaneous investigation of multiple traits. Such population-based cohorts are not ascertained by a particular clinical phenotype but instead can include information collated through linking national healthcare registries with data from questionnaires, multimodal imaging, physical measures, biomarker assays and genome-wide genotyping. The ability to query the impact of genomic variation on this wide range of phenotypes is set to lead to a step-change in the rate of genetic discovery, and such studies are already beginning to yield results. For example, genetic variants in the gene encoding the amyloid-β precursor protein have been identified that protect against Alzheimer’s disease in an Icelandic population cohort^8^, and a phenome-wide association study (PheWAS) of electronic health records from the eMERGE network has discovered effects of *IRF4* variants on actinic keratosis and non-melanoma skin cancer^9^.

However, capitalizing on the availability of population-based cohorts for biomedical research is complicated by the scale and nature of the data: the phenotypic space is multidimensional and heterogeneous, as data can be subject to observational predilections, nonuniform recording practices, and longitudinal biases, and the prevalence of phenotypic events is variable^10–15^. This creates new challenges that are not currently addressed by existing analytical methods for GWAS and PheWAS. An open question is how to achieve a balance in investigating the precise phenotypes obtainable from routine healthcare data at a level of resolution that has sufficient statistical power to detect genetic associations. For example, an assessment of a disease category, such as ankylosing spondylitis (AS), will be well powered if the disease in question is common in the population under investigation, but is unlikely to reveal informative associations above and beyond those identified through GWAS. Conversely, interrogating a diagnosis such as AS with cervical spine complications, could provide novel insight regarding the contribution of genetics to disease subphenotypes such as disease location or to comorbidities, but would be underpowered if the precise phenotype is rare.

An ideal analytical scenario would be to allow the data to determine the appropriate level of resolution for identifying genetic associations. In the context of gene expression analysis, for example, mixed models have been used to infer the low-rank correlation structure of phenotypes^16^ so as to improve the detection of genetic associations. However, for routine healthcare data from population-based cohorts, the phenotypic state space is too large and sparse for such explorations. Instead, making use of the hierarchical structure of many disease classifications, such as the tree of International Classification of Diseases, Tenth Revision (ICD-codes, provides a tractable solution. Here we have developed a novel Bayesian analysis framework for identifying genetic associations across the entire health phenotype space by taking advantage of the relative topology of nodes within two tree-structured phenotypic datasets from the UK Biobank - the self-reported (SR) diagnoses that are organised using the UK Biobank classification tree which includes 531 diagnostic terms, and the hospitalisation episode statistics (HES) data that utilise ICD-10 codes and contain 16,310 diagnostic terms.

## RESULTS

### Tree analysis approach

Given a collection of health records from a population cohort annotated with past and present diagnoses, we want to perform a test of association that determines if a genetic variant affects susceptibility to at least one of the clinical phenotypes observed in the UK Biobank. Furthermore, we want our statistical framework to meet a set of fundamental requirements. Firstly, the method must accommodate different types of genetic variation, such that association tests can be performed with (i) a single nucleotide polymorphism (SNP), (ii) a single haplotype in a highly polymorphic region like the human leukocyte antigen (HLA) gene region,or (iii) a genetic risk score (GRS) constructed using multiple SNPs or haplotypes, each known to be associated with a quantitative trait or a complex disease. Secondly, for single locus variation, the test of association must be able to accommodate any genetic model (e.g. additive, dominant or full). Thirdly, the method must allow for the joint analysis and quantification of the evidence of association at each of the individual clinical phenotypes, and must estimate the genetic coefficients of effects. Next, the method must allow the identification of independent genetic effects by performing conditional analysis. Lastly, the method must model the correlation structure of genetic effects across the observed clinical phenotypes using a priori knowledge of the relationship between phenotypes as obtained from a diagnosis classification tree.

Here we propose a novel Bayesian analysis framework, termed TreeWAS, which models the genetic coefficients across all phenotypes as a set of random variables. To model the correlation structure we allow the coefficients to evolve down a tree in a Markov process (**Fig. 1**). The tree structure is determined from a known classification hierarchy, where each node in the tree is a clinical term in the classification, and observations can be made at both terminal and internal nodes. The prior *θ* determines the expected degree of correlation between genetic coefficients across phenotypes. The coefficient at a parent node can either be inherited by a child node with probability *e^−θ^*, or can transition to a new uncorrelated value, with probability 1−*e^−θ^*. This new value will be zero with a probability 1−*π*_1_, or non-zero with a probability *π*_1_. Thus, the *θ* and *π*_1_ parameters define the transition probabilities that control the Markov process. Given the structure of the model and the Markov process assumption, we can calculate the likelihood over the genetic coefficients across all clinical phenotypes using dynamic programming (detailed derivation and implementation is provided in the **Supplementary Note**), and we estimate a Bayes Factor (BF) for the evidence that the genetic coefficients are non-zero for at least one node in the tree. We refer to this statistic as the BF_tree_. Similarly, because of the properties of the model, using dynamic programming and the forward and backward algorithms, we can also determine the marginal posterior probability (*PP*) at each node in the tree that the genetic coefficient is non-zero, and the magnitude of this effect using the maximum a posteriori (MAP) estimator (see **Supplementary Note** for details).

**Figure 1.**
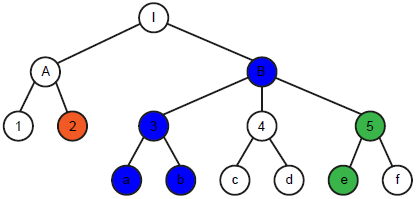
**Schematic of diagnosis classification tree and genetic coefficient transition scenarios tested**. Each node in the tree represents a clinical diagnosis and nodes are ordered in a hierarchical structure based on a classification criterion (such as similarities in clinical manifestations). White nodes represent the null state whereby there is no genetic association with the clinical phenotype. Green, red and blue nodes represent the alternative state whereby there is a genetic association with the clinical phenotype, with the different colours corresponding to different, uncorrelated genetic coefficients of association. A genetic coefficient can transition from the null state to a non-zero coefficient as in the I→ B and A→2 pairs. From the non-zero state a genetic coefficient can remain in a correlated non-zero state (as in the B→3, 3→a, 3→ b and 5→e pairs); it can transition back to the null state (as in the B→4 and 5→f pairs); or it can transition to a new, uncorrelated non-zero state (as in the B→5 pair). An in-depth description of the method is provided in the **Supplementary Note**.

### *HLA-B*27:05* TreeWAS and PheWAS comparison

We first illustrate the advantages of the TreeWAS approach, compared to existing PheWAS tests, by analysing the genetic association of the *HLA-B*27:05* allele against the UK Biobank HES dataset. The association of the *HLA-B*27:05* allele with AS is one of the strongest genetic effects observed in human complex diseases, with an estimated odds ratio of 46 (ref. ^17^), and this allele also confers risk for several other IMDs^18^, such as reactive arthritis^19^, psoriatic arthritis^20^ and anterior uveitis (iridocyclitis/iritis)^21^. Using PheWAS, where the evidence of genetic association for each observed clinical term is estimated independently, *HLA-B*27:05* is significantly associated with a total of six ICD-10 terms after correcting for multiple testing (*P-*adj < 0.05; using the Benjamini & Hochberg procedure^22^), including M45 AS and M45. X9 AS (Site unspecified) (**Fig. 2a**). However, this approach fails to identify associations with terms that have a greater granularity of clinical description and which are of a relatively low prevalence, such as M45.X6 AS with lumbar spine involvement (*P* = 0.01, *P*-adj = 1.0), which has a prevalence 17 times lower than the term M45. X9 (0.08%). By contrast, when employing the TreeWAS method, we observed *HLA-B*27:05* associations with 145 ICD-10 terms (*PP* ≥ 0.75), clustered in different branches of the classification tree (**Fig. 2b-e** and **Supplementary Table 1**). As for the PheWAS, there was a significant association with the M45 AS term (*PP* = 1), but the TreeWAS also revealed associations with four M45 subcategories all with the same estimated genetic effects (M45.X0, M45.X2, M45.X6 and M45.X9) rather than two (M45.X0 and M45.X9) (**Fig. 2a,b**). Moreover, there was evidence of association with the broader Spondylopathies category (M45-M49) (*PP* = 1.0), which was likely driven by the association with M45 (*PP* = 1.0), as well as M49 (*PP* = 0.43), but not with M47 Spondylosis (*PP* = 0.07) (**Fig. 2b**). Notably, as spondylosis is a non-immunological condition occurring as a consequence of age-related disk degeneration^23^, the lack of an *HLA-B*27:05* association with this node is consistent with an aetiology that is distinct to the autoimmune or infection-associated spondylopathies.

**Figure 2.**
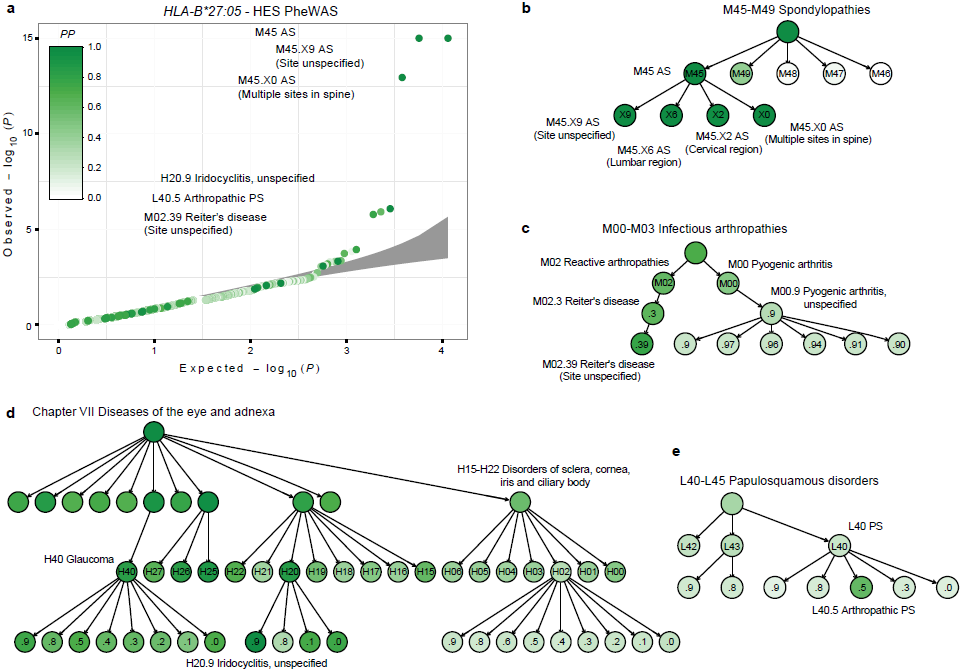
**Evidence of *HLA-B*27:05* allele association with risk for clinical diagnoses in the HES dataset. a**, Quantile-quantile plot of association test *P*-values of the *HLA-B*27:05* allele with each diagnosis term in the ICD-10 classification tree performed with maximum likelihood estimation using a logistic regression model. Grey area depicts the 95% confidence interval of sampling variance. Results are coloured-coded based on the posterior probability (*PP*) that *HLA-B*27:05* is associated with each diagnosis term as estimated with the TreeWAS model. **b-e**, Branches of the ICD-10 classification tree where significant associations between *HLA-B*27:05* and clinical diagnoses were identified (*PP* > 0.75). Results are tabulated in **Supplementary Table 1**. AS, ankylosing spondylitis; PS, psoriasis.

Associations with reactive arthritis (e.g. M02.39 Reiter’s disease; *PP* = 0.78) and anterior uveitis (H20.9 Iridocyclitis, unspecified; *PP* = 0.98) were observed as well (**Fig. 2c,d**). Interestingly, we also found a previously unreported *HLA-B*27:05* association with the term H40 Glaucoma (*PP* = 0.84) and related phenotypes (**Fig. 2d**); as glaucoma is a common complication of chronic uveitis^24^, comorbidity may explain this association. Lastly, we observed a weak effect on susceptibility to L40.5 Arthropathic psoriasis (PS) (*PP* = 0.60). Arthropathic PS develops in ~30% of PS patients, and is known to be associated with *HLA-B*27:05*, unlike non-arthropathic PS^18,25^ – which we confirmed through our approach (*PP* ≤ 0.25 for L40 child nodes other than L40.5) (**Fig. 2e**).

Our TreeWAS analysis of *HLA-B*27:05* in the HES dataset has recapitulated the known associations of this HLA allele. Moreover, we show how the TreeWAS method is capable of identifying additional bona fide associations compared to an approach where each ICD-10 term is considered independently, as in a PheWAS analysis.

### Sensitivity and specificity analysis of TreeWAS approach using simulated data

Given the greater capacity of TreeWAS to identify multiple associations with *HLA-B*27:05* we wanted to investigate the specificity and sensitivity of the method in greater depth. To assess the relative power of the TreeWAS method compared to alternative approaches for analysing routine healthcare data and to explore the robustness and accuracy of our method, we performed a set of simulations. In the first set, we assessed power by simulating data from a simple scenario where the genetic coefficients are non-zero for a set of five clinical annotations in the tree. These were chosen such that they were either found within a single branch of the classification tree, referred to as clustered nodes, or across distant branches of the tree, referred to as distributed nodes. We compared the power obtained under these two scenarios when considering a range of different allele frequencies. We fitted the TreeWAS model under a two-parameter setting with default parameters *θ* = 1/3 and *π*_1_ = 0.001, and for the alternative PheWAS model we assumed complete independence across annotations in the tree, equivalent to setting the parameter *θ → ∞*. Under the clustered nodes simulations, the relative gain in power for identifying active nodes, where the genetic coefficients are non-zero, of the TreeWAS model compared to the PheWAS model was 20-25% across the allele frequencies tested (**Fig. 3a**). This gain in power was not associated with an increase in the false positive rate (< 0.001), as observed in the nodes simulated with zero genetic coefficients (**Fig. 3a**). When we simulated non-zero genetic coefficients in distributed nodes we observed a 1-3% reduction in power to identify active nodes when using the TreeWAS method compared to the PheWAS approach (**Supplementary Fig. 2**). We also observed an increase in power (3.4-5.4%) in quantifying the evidence at the tree level with clustered nodes, but a small decrease in power with distributed nodes (0.2-1.0%) (**Supplementary Fig. 3 and 4**). These results illustrate that when the correlation in genetic coefficients is captured by the classification tree the gain in power with the TreeWAS method relative to the PheWAS approach is substantial, and if the correlation is not well-represented by the tree then the cost incurred by employing the former method is minimal.

**Figure 3.**
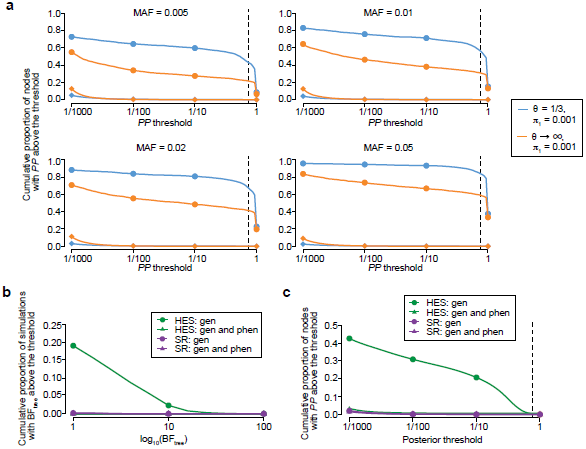
**Sensitivity and specificity analysis of TreeWAS on simulated data. a**, Rate of active node identification at increasing posterior probability (*PP*) thresholds and different simulated allele frequencies of the causal genetic variant, for the TreeWAS method (*θ*=1/3 and *π*_1_ =0.001) and a model assuming complete independence among phenotypes (*θ→∞* and *π*_1_=0.001; blue). For each simulation replicate we simulated five clustered nodes with non-zero genetic coefficients (•) and for the remaining nodes, phenotype counts were simulated to match observed disease prevalence and zero genetic coefficients (♦). Vertical dashed line denotes the *PP* threshold used in the analysis. Rate of false positives in the BF_tree_ statistic **(b)** and active node identification **(c)** when genotypes for the HLA-B*27:05 allele are permuted in both phenotypic datasets. Gen, genotype; phen, phenotype.

In the second set of simulations we assessed the impact of the non-independence between annotations arising from the way in which the clinical data are collected. For example, the recording of a specific disease subtype for any single individual may mean that other subtypes are less likely to be recorded for the same patient. We performed simulations under the null using the individual-level phenotype data from both of the UK Biobank phenotype datasets. For each simulation we permuted the observed genotypes of the *HLA-B*27:05* allele, representative of a common genetic variant (as its allele frequency is 4.05% in the UK Biobank), whilst maintaining the observed non-independence between annotations in the tree. For comparison, we also performed permutations of the individual-level phenotype data in addition to the genetic data, where all correlation is removed. With these permutations we quantified the rate of false positives in our analytical approach. When we permuted genotypes only, we observed an inflation of the BF_tree_ statistic and the node-level *PP* with the HES dataset, consistent with the more prominent correlation structure in the ICD-10 diagnosis tree compared to the SR diagnosis tree (**Fig. 3b,c**). Through these simulations we estimated a false positive rate of 0.05 and 0.01 with a log_10_ BF_tree_ threshold of 10 and 20, respectively, in the HES dataset, when substantial non-independence between nodes exists. For the SR dataset, the false positive rate at these thresholds was below 0.01. Thus, although non-independence between nodes can artificially increase the test statistics, we show that this can be countered with the use of conservative levels of significance such that the false positive rate can be maintained at an appropriate level.

### The effects on HLA allelic variation in the phenome

Genetic variation in the HLA region is associated with a wide range of human disorders, in particular autoimmune and autoinflammatory diseases, and thus we sought to interrogate the effect of HLA on the full range of SR and HES phenotypes using our TreeWAS approach. Through conditional analysis (Online Methods and **Supplementary Note**), we identified independent associations for ten HLA alleles in the SR data (log_10_ BF_tree_ ≥ 10) and eight in the HES data (log_10_ BF_tree_ ≥ 20) (**Fig. 4** and **Supplementary Tables 2** and **4**). Seven of these HLA alleles or alleles in high linkage disequilibrium (LD; *r* > 0.98) were found to be associated in both datasets (**Supplementary Fig. 5**).

**Figure 4.**
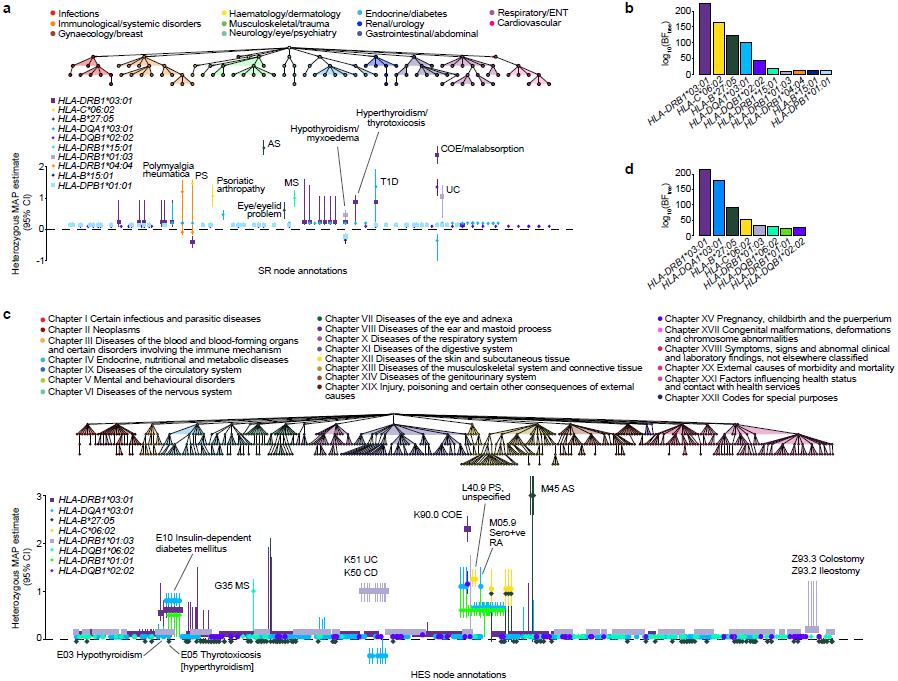
**Genetic analysis of HLA allelic variation in the risk of clinical phenotypes from the UK Biobank SR diagnosis and HES datasets. a**, The tree depicts the hierarchical structure of self-reported clinical phenotypes as determined by the UK Biobank classification. Only nodes with a significant association (*PP* > 0.75) with at least one HLA allele are shown, along with their parent nodes. The graph shows estimated effect sizes for the heterozygous genotype of the different HLA alleles on susceptibility to each clinical phenotype. Bars show the 95% credible interval. **b**, Evidence of association for each HLA allele with at least one node in the tree (BF_tree_) in the conditional TreeWAS analysis for the SR dataset. **c**, The tree depicts the hierarchical structure of HES-derived clinical phenotypes as determined by the ICD-10 classification (showing nodes with *PP* > 0.75 and their parent nodes). The graph shows estimated effect sizes for the heterozygous genotype of the different HLA alleles on susceptibility to each clinical phenotype. **d**, Evidence of association for each HLA allele with at least one node in the tree in the conditional TreeWAS analysis using the HES data. Estimates for heterozygous and homozygous genotype effect sizes and descriptions of all phenotypes shown are available in **Supplementary Tables 2** and **4**. AS, ankylosing spondylitis; CI, confidence interval; COE, coeliac disease; ENT, ear, nose, throat; MAP, maximum a posteriori; MS, multiple sclerosis; PS, psoriasis; RA, rheumatoid arthritis; T1D, type 1 diabetes; UC, ulcerative colitis.

**Figure 5.**
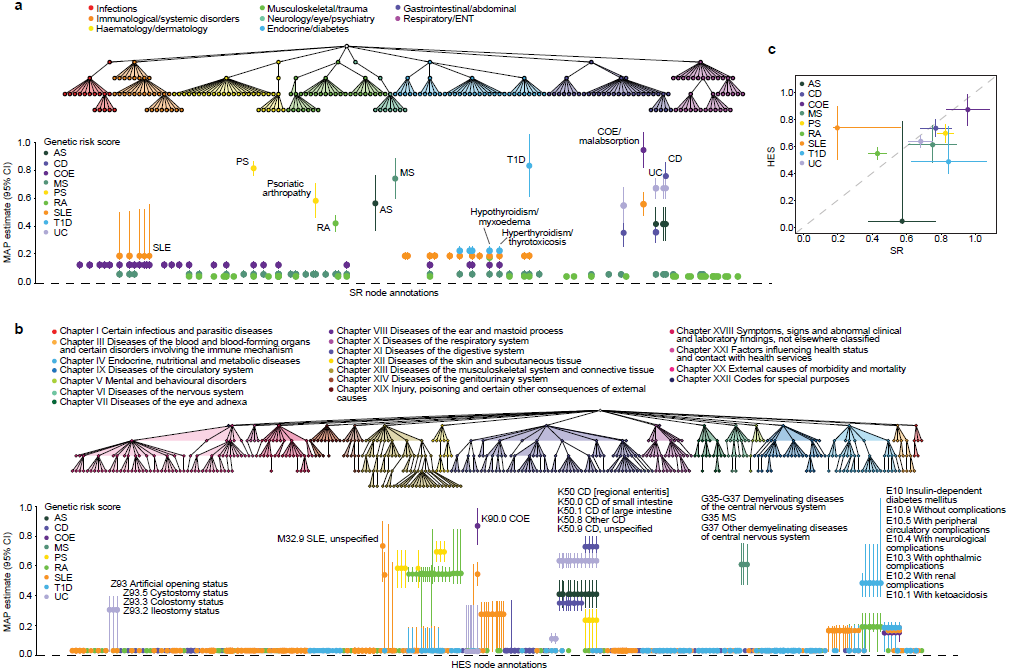
**Association analysis of genetic risk for multiple IMDs derived from clinical phenotypes in the UK Biobank SR diagnosis and HES datasets. a**, The tree depicts the hierarchical structure of SR clinical phenotypes as determined by the UK Biobank classification. Only nodes with a significant association (posterior probability > 0.75) with at least one IMD genetic risk score (GRS) are shown, along with their parent nodes. The graph shows estimated effect size of GRS on susceptibility to each clinical phenotype with posterior probability > 0.75. Bars show the 95% credible interval. **b**, The tree depicts the hierarchical structure of HES–derived clinical phenotypes as determined by the ICD-10 classification (showing nodes with posterior probability > 0.75 and their parent nodes). The graph shows estimated effect sizes of GRS on susceptibility to each clinical phenotype. **c**, Comparison of estimated genetic coefficients for each GRS and the respective clinical annotation in both phenotypic datasets. Estimates of effect sizes and description of all phenotypes shown are available in **Supplementary Table 8**. AS, ankylosing spondylitis; CD, Crohn’s disease; CI, confidence interval; COE, coeliac disease; ENT, ear, nose, throat; MAP, maximum a posteriori; MS, multiple sclerosis; PS, psoriasis; RA, rheumatoid arthritis; SLE, systemic lupus erythematosus; T1D, type 1 diabetes; UC, ulcerative colitis; MAP.

These associations were fine-mapped, and the majority of the strongest effects were with IMDs as reported previously through^17, 26–31^ GWAS (**Fig. 4**). For the class I alleles, we observed associations with PS (*HLA-C*06:02*) and AS (*HLA-B*27:05*), with the genetic coefficients of the latter being the largest observed in both the SR and HES datasets (**Fig. 4a,c**). For the class II alleles, *HLA-DRB1*03:01* and *HLA-DQB1*02:02* were observed to be independently associated with coeliac disease (COE) in both datasets; these alleles tag two of the strongest known COE HLA risk haplotypes, DR3-DQ2 and DR7-DQ2 (ref. ^27^). In both datasets, *HLA-DQA1*03:01* was identified and fine-mapped to rheumatoid arthritis (RA); this allele is in moderate LD with *HLA-DRB1*04:01*(*r* = 0.71), which is the likely causal allele driving this association^28^. Similarly, *HLA-DQA1*03:01* was also found to be associated with type 1 diabetes (T1D), noting that this allele is in LD with *HLA-DQB1*03:02 (r* = 0.67), which has been indicated as the most significantly associated class II allele in T1D^26^. In the SR dataset we identified an *HLA-DRB1*15:01* association and fine-mapped it to multiple sclerosis (MS) (**Fig. 4a**). In the HES dataset *HLA-DQB1*06:02* was identified instead and also fine-mapped to MS (*PP* = 1; **Fig. 4c**), but this allele is in strong LD with *HLA-DRB1*15:01* (*r* = 0.97) (**Supplementary Fig. 6**). Lastly, *HLA-DRB1*01:03* was fine-mapped to ulcerative colitis (UC) and Crohn’s disease (CD) in both datasets, and this allele has been previously found to be the likely causal allele for these two types of inflammatory bowel disease (IBD)^31^.

Apart from established HLA associations with common IMDs, we also confirmed HLA effects for conditions where GWAS have not been performed, we detected associations with clinical annotations linked to disease complications, and we identified novel HLA associations with other IMDs. For example, in the SR dataset, we confirmed the association of *HLA-DRB1*04:04* with polymyalgia rheumatic and giant cell arteritis, which has been previously identified only through small candidate gene studies^32, 33^ (**Fig. 4a**). The UC- and CD-associated *HLA-DRB1*01:03* allele was found to also be associated with surgical procedures linked to complications of IBD, such as Z93.3 Colonoscopy status (*PP* = 1) and Z93.2 Ileostomy status (*PP* = 1), consistent with findings by the International IBD Genetics Consortium^34^ (**Fig. 4c**). Of the ten HLA alleles independently associated with clinical phenotypes in the SR dataset, five were associated with hypothyroidism/myxoedema, and three of the eight HLA alleles from the HES data were associated with the E03 hypothyroidism code - thereby constituting this disease as the phenotype with the largest number of independent HLA association across both UK Biobank datasets. Associations have been reported with hypothyroidism for both the HLA class I and class II loci, but the specific HLA alleles driving these have not been well resolved^35, 36^, apart from a recently reported *HLA-DQA1*05:01-HLA-DQB1*02:01-HLA-DRB1*03:01* (HLA-DR3-DQ2 haplotype) association^37^. Further to *HLA-DRB1*03:01*, we have refined the HLA associations with hypothyroidism to two additional independent risk alleles, *HLA-DQA1*03:01* and *HLA-DRB1*01:03*, and two independent protective alleles, *HLA-B*15:01* and *HLA-DPB1*01:01*(**Fig. 4** and **Supplementary Table 10**). Our HLA analysis of the UK Biobank routine healthcare data therefore demonstrates the validity of our method in that it can identify known genetic associations, and can additionally facilitate the discovery of new associations for diagnoses that have as yet been relatively understudied.

### Genetic risk score associations with IMDs

Outside of the HLA, over the last decade our understanding of genetic susceptibility to the common IMDs has increased tremendously as tens to hundreds of risk loci have been identified per disease^38^. We computed GRSs, which aggregate the genetic effect of multiple susceptibility variants, for nine common autoimmune and auto-inflammatory diseases (see Online Methods), and assessed their relationship with the phenome in the UK Biobank (**Fig. 5**).

In most cases the GRSs best identified those clinical annotations from which they were constructed, with secondary associations being detected for conditions with shared genetic risk. For example, CD and UC have a high genetic correlation^39^, although disease-specific susceptibility loci have been identified for each and heterogeneity in effect sizes has been observed^40^. The GRS for CD was thus associated with both CD itself as well as UC, but the magnitude of genetic coefficients was greater for CD as expected (*β*= 0.86 vs. *β*= 0.44 in SR and *β*= 0.73 vs. *β*= 0.35 in HES for CD and UC, respectively). However, the GRS for UC could not differentiate these two clinical annotations, with estimated genetic coefficients of the same magnitude for both CD and UC (*β*= 0.68in SR and *β*= 0.64 in HES), indicating some level of variation in the precision of different GRSs to identify specific phenotypes (**Fig. 5a,b**).

For all associations, including those of GRSs with their respective clinical annotations, the genetic coefficients were less than 1, demonstrating a degree of dilution in phenotype detection across both the SR and HES datasets. The least dilution was observed for the association of the GRS for COE and this disease (*β*= 0.96 and *β*= 0.87 in the SR and HES datasets, respectively). This suggests that the COE phenotypes derived from the UK Biobank routine healthcare data are highly comparable to the clinically ascertained disease phenotype used in the GWAS^41^ from which the variants for the COE GRS were obtained. Across both datasets the greatest dilution of a GRS and its respective disease was observed for RA (*β*= 0.43 and *β*= 0.55 in the SR and HES data, respectively), whilst in the HES data specifically the AS GRS was not associated with the disease (*PP* = 0.01) potentially due to the small number of AS patients in this dataset (*n* = 146), and in the SR data the SLE GRS association with this disease had a genetic coefficient of only 0.20 (**Fig. 5a,b**).

Overall the GRS associations observed were largely consistent across the SR and HES datasets, and for the GRSs and their respective diseases, the estimated genetic coefficients were weakly positively correlated (*r*_corrected_ = 0.23; correcting for measurement error) across both datasets (**Fig. 5c**). Strikingly, although the capacity of the SLE GRS to identify SLE itself in the SR data was so low that the SLE GRS was in fact a better predictor of COE (*β* = 0.57) (**Fig. 5a**), in the HES dataset this was not the case. The SLE GRS was most predictive of the M32.9 SLE term (*β* = 0.50; *PP* = 1.00), and to a lesser extent of K90.0COE (*β* = 0.47; *PP* = 1.00) (**Fig. 5b**). This discrepancy between the SR and HES datasets with respect to these findings suggests differences in the diseases annotated as SLE in the two datasets, which may in turn reflect differences in disease perception or diagnosis that could have clinical implications. Notably, in the SR data the SLE node was also associated with the COE GRS (*β* = 0.13), but this was not the case in the HES data, further supporting a distinction between the SLE phenotypes in the two datasets.

Secondary associations of the GRSs were identified either with complications that were corollaries of the disease with which the primary association was observed, or with other IMDs. For example, as observed for the associations with the *HLA-DRB1*01:03* allele, the GRS for UC was associated with colostomy and ileostomy events (*β* = 0.31 and *PP* = 0.98, and *β* = 0.31 and *PP* = 1, respectively), and we observed the same associations with the CD GRS, although the magnitude of the effect sizes was lower (*β* = 0.03 and *PP* = 0.91, and *β* = 0.03 and *PP* = 0.87, respectively). Also paralleling the HLA analysis, hypothyroidism was found to be associated with several GRSs in the SR and HES datasets. We found that five and four of the nine GRS tested were associated with hypothyroidism in the SR and HES datasets, respectively, with those for COE, RA, SLE and T1D being found in both datasets. Thus, hypothyroidism is the single phenotype with the largest number of different GRS associations (**Fig. 5a,b** and **Supplementary Table 6 and 8**).

## DISCUSSION

Our Bayesian analysis framework addresses a fundamental challenge for the analysis of high-dimensional, heterogeneous routine healthcare data - chiefly how to identify statistically significant genetic associations when interrogating the thousands of diagnoses included in these data. Rather than using phenotype aggregation or employing traditional methods of dimension reduction that would sacrifice phenotypic resolution^10, 12^, we have instead exploited the inherent hierarchical structure of diagnostic classifications to investigate the impact of genetic variation across the full breadth of the UK Biobank phenome. Our TreeWAS approach allows the calculation of well-calibrated statistics for quantifying the evidence of genetic association, and through simulations we demonstrate that compared to PheWAS, the inference of genetic coefficients by TreeWAS is improved by over 20% when the hierarchical structure of phenotypic nodes correctly represents the genetic relationship between phenotypes. Thus, when applying the TreeWAS method to interrogate the effect of HLA on the UK Biobank phenome, associations were identified with 143 and 966 nodes in the SR and HES datasets, respectively. Assessing the impact of GRSs derived from clinically confirmed IMD cohorts also revealed associations with 151 and 810 nodes in the two respective healthcare datasets. The total number of nodes identified demonstrates the power of the TreeWAS approach in detecting associations in datasets where numerous weak but correlated effects are present across the classification tree.

Amongst the many active nodes for which genetic associations were observed, previously established effects of HLA alleles on specific IMDs were detectable, as were effects for relatively understudied conditions. Notably, multiple novel associations with HLA class I and II alleles were discovered for hypothyroidism, including effects conferring risk for and protection against this disease. Although not all previously reported HLA associations could be detected for any single IMD - such as AS^42^ or MS^30^ - due to limited power with the current UK Biobank datasets, the capacity for genetic discovery will improve with increasing cohort size, and associations with nodes displaying a substantial granularity of clinical description were already identifiable. For example, *HLA-B*27:05* was associated not only with M45 AS but also with subphenotypes of this disease node which specify the spinal locations where the pathology occurs, and *HLA-DRB1*01:03* was associated with the CD and UC nodes as well as with surgical procedures linked to these conditions.

In the GRS analysis, associations between GWAS-derived GRSs and their respective diseases were typically the strongest effects observed, even in the absence of HLA allele inclusion, hence demonstrating that non-HLA variants can provide precision for detecting specific IMDs. Cross-disease associations of GRSs were also identified; in particular, hypothyroidism was associated with the largest number of different IMD GRSs in both the SR and HES datasets. This extent of genetic sharing has not been previously appreciated and indicates a common, genetically determined aetiopathogenesis. For all GRS associations, dilution of the capacity for phenotype detection was observed, as all genetic coefficients were below 1. The degree of dilution varied across different diseases, but was largely comparable between the SR and HES datasets. An intriguing exception, however, was the differential association of the SLE GRS with the respective SLE terms in the SR and HES data, such that this GRS could not precisely predict the self-reported disease, but could accurately detect the hospitalisation record-derived phenotype. Compared to the other IMDs for which GRSs were constructed, SLE is a more heterogeneous condition in which various different organs may be affected and which consequently presents a substantial diagnostic challenge^43^. Therefore, this discrepancy in the magnitude of the SLE GRS associations could reflect inaccurate reporting of the disease by UK Biobank participants, an over-diagnosis of the disease that is not discernible from the HES data if hospitalisation is associated with a more clear-cut diagnosis, or a greater disease heterogeneity whereby SLE as defined in GWAS and in the HES data represents only a subset of a more genetically variable syndrome. This example demonstrates how exploring the genetic basis of the healthcare phenome, particularly when incorporating the use of different phenotypic datasets, can help to expose disease areas where improvements are required to ameliorate disease perception or strengthen diagnostic practices, for instance.

Our TreeWAS method provides a robust framework for investigating the impact of genetic variation on the complete range of phenotypes annotated within routine healthcare data. We have used disease classification trees to model the correlation between genetic effects, and an extension of our current method could include the modelling of the parameters *θ* and *π*_1_, by incorporating a prior distribution or by estimation through inference methods, to permit different transition rates to be interrogated along the tree for improved modelling of the genetic relationship between nodes. Our method could also be further developed so as to use the data to learn about the correlation structure - allowing for less restricted topologies in disease classifications - moving from a tree structure to a generalised directed acyclic graph. The generation and modification of approaches such as TreeWAS are of particular importance as the entire space of the human phenome expands. The increased incorporation of correlated, high-dimensional phenotypes, such as measures of temporal disease progression^44^, is anticipated and their wider analysis will necessitate methodological evolution to combine the multiple dimensions of these complex phenotypes in order to provide an integrated outcome.

**URLs.** UK Biobank, http://www.ukbiobank.ac.uk/; UK Biobank genotyping procedure and genotype calling protocols, http://biobank.ctsu.ox.ac.uk/crystal/refer.cgi?id=155580; UK Biobank internal quality control procedures, http://biobank.ctsu.ox.ac.uk/crystal/refer.cgi?id=155580; HLA*IMP, https://oxfordhla.well.ox.ac.uk/hla/; World Health Organisation ICD-10 disease classification codes, http://www.who.int/classifications/icd/en/.

## ONLINE METHODS

### UK Biobank data

The UK Biobank is a prospective cohort of over 500,000 men and women aged 40 to 69 years when recruited in 2006-2010. Participants have provided: data on lifestyle, environment, and medical history through an interview and completion of a questionnaire; physical measures; biological samples for genotyping and biochemical assays; and informed consent to long-term medical follow-up through linkage of national health registries. UK Biobank data is available under open access to conduct health-related research after approval of a project proposal^6^. The UK Biobank has obtained ethical approval covering this study from the National Research Ethics Committee (REC reference 11/NW/0382).

### Phenotypic data

We analysed two phenotypic datasets available through the UK Biobank. The first included the SR diagnosis data, ascertained through the completion of questionnaires and interviews with study participants (data field 20002 Non-cancer illness code,self-reported); the second dataset included the HES registry dataset ascertained through linkage of health registries (data fields 41142 and 41078; accessed on September 2016). Clinical diagnoses in these datasets are described with different classification schemes, both of which follow a hierarchical structure. The diagnosis terms used to store the medical history of UK Biobank participants were proposed by the UK Biobank team (data-coding 6), and this classification tree is organised into 11 subclasses with a total of 561 clinical terms, 531 of which are selectable. Diagnosis terms used to store hospitalisation events follow the ICD-10 list compiled by the World Health Organisation. The ICD-10 classification tree is organised into 22 Chapters and containing a total of 19,855 clinical terms, 16,310 of which are selectable. Each hospitalisation episode in the dataset has a primary diagnosis associated with the event and an event may be annotated with one or more secondary diagnoses. Disease outcomes for each individual, as a binary trait, were generated for the combined primary and secondary diagnoses annotations.

### Genetic dataset

The interim release of the UK Biobank genetic data used for this study includes 152,732 individuals, 120,286 of which were determined to be of British Isles ancestry (**Supplementary Fig. 6**). The initial 50,000 individuals were genotyped on the Affymetrix UK BiLEVE Axiom array as part of a pilot study described elsewhere^45^ and the remaining 102,732 individuals were genotyped on the Affymetrix UK Biobank Axiom array. The quality control of SNP data and whole-genome SNP imputation was performed by the UK Biobank analysis team and described in the UK Biobank website (http://www.ukbiobank.ac.uk/scientists-3/genetic-data). We imputed 356 classical HLA alleles for the *HLA-A*, *-B*, *-C*, -*DRB5*, -*DRB4*, -*DRB3*, -*DRB1*, -*DQB1*, -*DQA1*, -*DPB1*and -*DPA1* loci at four digit resolution with the HLA*IMP:02 algorithm^46^ using data from a multi-population reference panel. The imputation panel contained 2,263 SNPs in the MHC region (GRCh37 coordinates chr6:29500000-33500000) which overlapped UK Biobank genotyped SNPs. This SNP set was selected to optimize MHC coverage and imputation performance and the HLA*IMP:02 algorithm was trained on this SNP set. Genetic risk scores, weighted by effect sizes, were generated for nine IMDs using genome-wide associated variants compiled from previous studies: AS^17^, CD^40^, COE^41^, MS^47^, PS^26^, rheumatoid arthritis^48^, SLE^49^, T1D^50^, and UC^40^. SNP genotypes for the UK Biobank individuals were extracted from the imputed genotype data.

### Simulated data

To assess the accuracy of the method, we simulated case-control status for 120,000 individuals and the 531 selectable phenotypes in the diagnosis tree used for the self-reported dataset and with disease prevalence as observed in the UK Biobank cohort. Simulations were generated under two scenarios. For the first, we assumed a causal relationship between a genetic variant and five clinical terms under the same parent node in the tree (disease prevalence in these nodes ranged between 0.01 and 0.4%). These simulations are referred to as clustered clinical phenotypes. The second set of simulations, termed distributed phenotypes, consisted of five clinical terms with a causal relationship distributed under different branches of the classification tree; these clinical terms were selected with matching disease prevalence, as for the clustered simulations. For each scenario we simulated genotypes sampled from a multinomial distribution with a fixed allele frequency and genetic coefficients sampled from the prior (**Supplementary Figure 1**). Case-control status was determined by using logistic risk with a y-intercept matching the observed disease prevalence. Sets of simulations were performed for the allele frequencies 0.005, 0.01, 0.02 and 0.05. For each simulation we computed the evidence of association in the tree (BF_tree_), and the evidence of association at each individual node with the parameters *θ* = 1/3 and *π*_1_ = 0.001. We compared the power to detect association with at least one node in the tree with an analysis where we assume no correlation in the genetic coefficients between nodes in the tree, equivalent to setting *θ ⟶ ∞* in the TreeWAS method (see **Supplementary Note**). 500 simulation replicates were performed for each combination of parameters and settings. To assess the robustness of the algorithm to the non-independence between annotations unaccounted by the tree structure, we performed simulations where we permuted the genotypes whilst leaving the observed phenotypes in the UK Biobank cohort intact. Simulations were performed with the observed self-reported and HES datasets, and we permuted the observed genotype.

### HLA analysis

Best-guess genotypes were used to generate count distributions in affected and unaffected individuals at each terminal node in the tree. To identify independent HLA associations we performed sequential conditional analysis using an approximation to the likelihood function as described in the **Supplementary Note**. At each step, BF_tree_ statistics were generated for each allele and the allele with the largest was selected for conditioning in the next iteration. Conditional analysis was repeated until all observed BF_tree_ statistics were below 10^10^ in the self-reported diagnosis dataset and 10^20^ in the HES dataset, ensuring a false discovery rate below 0.01, as determined through the simulation analysis. For each significant allele association we computed the marginal *PP* for the genetic coefficient being not equal to 0 and the MAP estimate using posterior decoding as described in the **Supplementary Note**. Association with a clinical annotation was deemed significant if the *PP* was above 0.75.

## ACKNOWLEDGEMENTS

This research has been conducted using the UK Biobank Resource (application number 10625). The research has been supported by the Wellcome Trust (100956/Z/13/Z and 090532/Z/09/Z to G.M. and 100308/Z/12/Z to L.F.) and the Danish National Research Foundation (L.F.). This work was supported by the Australian National Health and Medical Research Council (NHMRC), Career Development Fellowship ID 1053756 (S.L.); and by the Victorian Life Sciences Computation Initiative (VLSCI) grant number VR0240 on its Peak Computing Facility at the University of Melbourne, an initiative of the Victorian Government, Australia (S.L.). Research at the Murdoch Childrens Research Institute was supported by the Victorian Government’s Operational Infrastructure Support Program.

## AUTHOR CONTRIBUTIONS

A.C. and G.M. performed the analyses with contributions from C.A.D. A.C., C.A.D., L.J., P.D., L.F. and G.M. conceived the study. HLA imputation was performed by A.M., D.V., A.D. and S.L. A.C., C.A.D. and G.M. wrote the manuscript and all other authors reviewed the manuscript.

